# Bisphosphonates Trigger Anti-Ageing Effects Across Multiple Cell Types and Protect Against Senescence

**DOI:** 10.1101/2025.03.25.645228

**Authors:** Jinsen Lu, Srinivasa Rao Rao, Helen Knowles, Haoqun Zhan, Beatriz Gamez, Eleanor Platt, Lucy R. Frost, Tiffany-Jayne Allen, Gayle Marshall, Kilian V.M. Huber, Ludwig G. Bauer, Iolanda Vendrell, Benedikt Kessler, Anne Horne, Ian R Reid, Chas Bountra, James L Kirkland, Sundeep Khosla, F Hal Ebetino, Emilio Roldan, R Graham G Russell, James R Edwards

## Abstract

Bisphosphonates (BPs) have been the major class of medicines used to treat disorders of excessive bone loss for over five decades. Recently it has been recognized that BPs may also have additional significant beneficial extra-skeletal effects. These include a reduction of all-cause mortality and of conditions commonly linked to ageing, such as cancer and cardiovascular disease. Here we show that bisphosphonates co-localize with lysosomal and endosomal organelles in non-skeletal cells and stimulate cell growth at low doses. *In vivo* spatial transcriptomic analysis revealed differentially expressed senescence markers in multiple organs of aged BP-treated mice, and a shift in cellular composition toward those of young counterparts. Similarly, a 5000-plex plasma proteome analysis from osteopenic patients before and after BP-treatment showed significant alterations in ∼400 proteins including GTPase regulators and markers of senescence, autophagy, apoptosis, and inflammatory responses. Furthermore, treatment with BPs protected against the onset of senescence *in vitro*. Proteome-wide target deconvolution using 2D thermal profiling revealed novel BP-binding targets (PHB2, ASAH1), and combined with RNA- and ATAC-seq of BP-treated cells and patient data, suggests downstream regulation of the MEF2A transcription factor within the heart. Collectively, these results indicate how BPs may beneficially modify the human plasma proteome, and directly impact multiple non-skeletal cell types through previously unidentified proteins, thereby influencing a range of pathways related to senescence and ageing.

## Introduction

Ageing is associated with the development of multiple morbidities. Common factors exist linking the control of lifespan with the development of age-related disorders (e.g. cardiovascular disease, neurodegeneration, bone loss). Moreover, multiple diseases of ageing can be simultaneously underpinned by dysregulation in a single unitary ageing mechanism, such as senescence, autophagy, oxidative stress management, or accumulation of DNA damage^1,2^. A single intervention may therefore have the potential to impact multiple disorders simultaneously and prevent occurrence of many others.

The bisphosphonate (BP) class of drugs has been used worldwide as an effective treatment for disorders of excessive bone loss (e.g. osteoporosis, cancer-induced osteolysis) over the past 50 years. BPs have a unique ability to bind to calcified surfaces which, coupled with their ability to inhibit bone resorption through cellular mechanisms, has led to several different BPs being developed for clinical use^3^. The earliest BPs to be used were clodronate (CLO), etidronate (ETI). Biochemically they at through different mechanisms than the more recently developed nitrogen-containing BPs (N-BPs), such as pamidronate (PAM), alendronate (ALN), risedronate (RIS), ibandronate (IBN), and notably zoledronate (ZOL)^4^. The bone-resorbing osteoclast is the primary bone cell capable of degrading and removing large quantities of bone. The ingestion of bone-bound N-BPs triggers an inhibition of farnesyl pyrophosphate synthase (FPPS), a key enzyme in the mevalonate pathway governing cholesterol biosynthesis. Blockade of the crucial intracellular mevalonate metabolic pathway impairs the prenylation of multiple GTPases, alters intracellular signaling events, and induces osteoclast dysfunction and apoptosis^5,6^.

More recently, studies of patients receiving BPs for skeletal-related conditions have revealed a reduced mortality^7–12^ and a reduction in the development of a number of disorders commonly associated with ageing, such as cardiovascular and respiratory disorders, cancer, along with improved overall survival following admission to intensive care units^5–8^. Such beneficial effects cannot be accounted for by the prevention of bone loss alone, challenging the bone- and osteoclast-specific nature of BP activity. The mechanisms underlying these effects remain largely unknown, indicating that the pleiotropic pharmacology of BP action remains incompletely understood and potential polypharmacology of BPs untested. This study was designed to explore the potential causative mechanisms of the non-skeletal “beyond bone” effects of BPs *in vivo*, using an integrated multiomics approach, including protein-binding, as well as alterations to the human secretome, transcriptional control and cellular impact.

## Results

### Bisphosphonate treatment reduces risk of disease

A series of bone loss-related randomized controlled trials^9–23^ across the globe suggests an overall trend of BPs to reduce mortality, with RIS the most frequently tested (35.6%), and ZOL (23.4%) showing the greatest significant benefit, along with Ibandronate (23.8%) and ALN (17.3%), in primarily female cohorts across all continents (**Fig.1a**). Furthermore, a systematic evaluation of BP treatment (clinical and pre-clinical)^13–80^ revealed significant associations with improved outcomes for diseases at multiple organ and tissue sites (**Fig.1b**). This included cardiovascular disease (e.g. vascular calcification, stroke, atherosclerosis, myocardial infarction), cancers (e.g. oesophageal, colorectal, prostate, breast, multiple myeloma, osteosarcoma, cervical, and ovarian), diabetes, neurodegenerative disorders, hearing loss, hereditary diseases such as Pseudoxanthoma Elasticum (PXE), Hutchinson-Gilford progeria syndrome, parasitic infections (Chagas disease, leishmaniasis), and rheumatic diseases (ankylosing spondylitis, rheumatoid arthritis).

**Figure 1.**
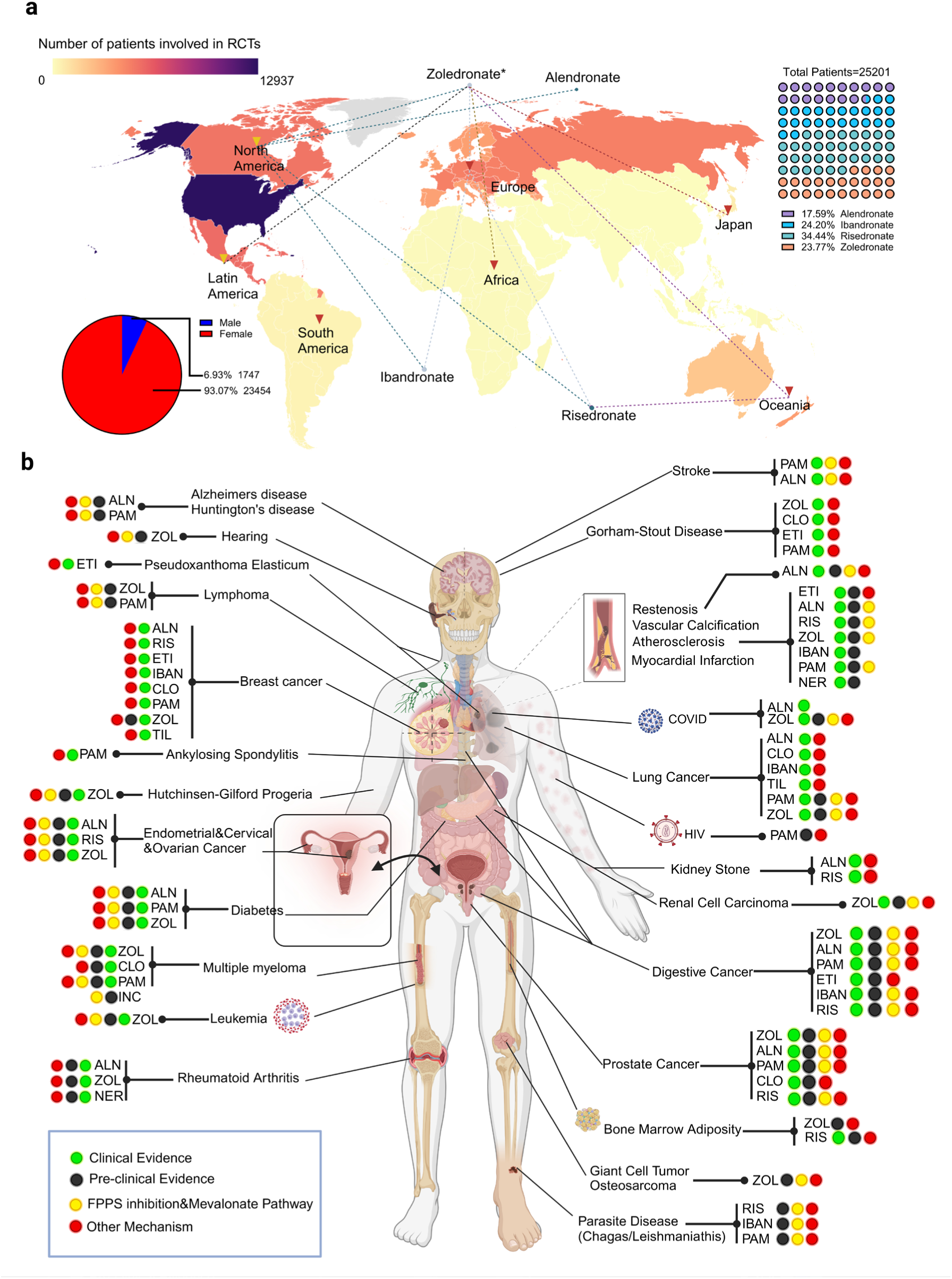
Clinical and preclinical evidence of bisphosphonate benefit. **a**. Geographical heatmap depicting global distribution of RCTs related to mortality reduction conferred by BPs. Heatmap = number of patients in a specific region. Colour = BP. RCT regions labelled and linked by dotted line. Gender distribution of participants patients illustrated as pie chart. **b**. Body based illustration of BP benefit against disease. Green=clinical study, Black=pre-clinical study, Yellow= benefit attributed to FPPS/mevalonate pathway inhibition, Red=other mechanism. PAM: Pamidronate; IBA: Ibandronate; TIL: Tiludronate; NER: Neridronate; INC: Incadronate.*p<0.05 vs placebo group (mortality rate reduction).

### Zoledronate treatment alters the human proteome

Plasma samples collected from patients^81^ before and after receiving ZOL (18 and 36 months) were assessed for changes in systemic protein levels (SomaScan, 5000-plex, SomaLogic). A significant number of proteins (352, log_2_FC>0.5, padj< 0.05) were shown to be differentially expressed 18 and 36 months following treatment with ZOL (**Fig.2a**). The highest regulated individual proteins included KLC1, FER, TMOD2, FYN, PPFIA1, CLINT1, SMTN, TANK, AMPD2, and LTB4R (**Fig.2a**), while GO based pathway analysis indicated altered expression of protein groups governing immune cell apoptosis, autophagy, oxidative stress response, telomere regulation, and endosome and lysosome organization, along with those linked to bone turnover (**Fig.2b**). Similarly, a substantial fraction of differentially expressed proteins correlated with ageing-related genomic signatures in the Human Ageing Genomic Resources (HAGR) database (**Fig.2c**) including markers of longevity, diet-restriction stimulated lifespan extension, and control of cellular senescence. Furthermore, 19 differentially expressed proteins were identified as common members of the toxic cocktail released by senescent cells known as the senescence-associated secretory phenotype (SASP) (including downregulation of inflammatory mediators IL-6, NFĸB), suggesting a seno-modifying potential for ZOL (**Fig.2d**).

**Figure 2.**
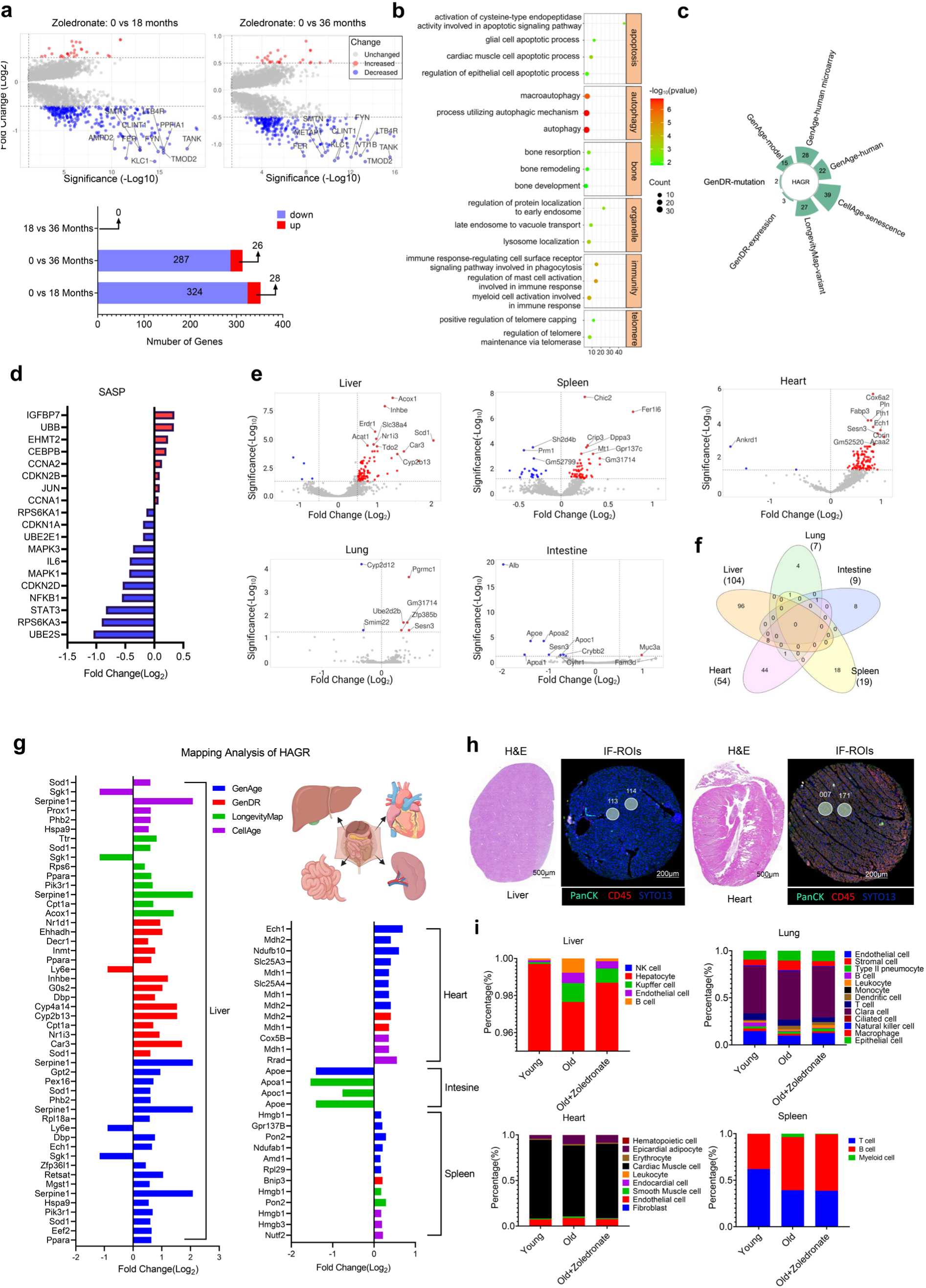
Zoledronate alters human plasma proteome and local organ transcriptome in ageing mice. **a**. Differential analysis of plasma protein alterations following ZOL treatment. Volcano plot illustrating DEPs between 0 and 18 mth, 0 and 36 mth groups. Bar chart indicating DEP number/group. Filter standard: |log_2_FC| >0.5 and padj<0.05. **b.** Enrichment analysis indicating ageing-related biological process (GO) in plasma DEPs (0 vs 18 months) categorized as apoptosis, autophagy, bone, endosome/lysosome, immune response, and telomere. **c.** Mapped DEPs in HAGR (CellAge, GenAge, GenDR, Longevity Map. **d**. Altered SASP in plasma DEPs (padj <0.05), mapped to SenoMayo panel, illustrated according to log_2_FC. **e,f.** DEGs in heart, liver, spleen, lung, and intestine of ageing mice (vehicle vs ZOL). **g**. DEGs mapped to HAGR databases (log_2_FC). **h**. Organ-specific ROIs selected from IF (PanCK, CD45, SYTO13) stained representative sections (eg. liver, heart), enumerated prior to spatial transcriptomic profiling**. i**. Organ-specific cell population deconvolution analysis (young vs aged, vehicle vs ZOL) using Tabula Muris database, CIBERSORTx comparing cell type proportions across liver, heart, spleen, intestine, and lung tissues.

### Zoledronate induces organ-specific transcriptomic ageing-related changes in vivo

The significant shift in the plasma proteome reflects a systemic ZOL-induced alteration within the blood compartment of female osteopenic patients but on its own, does not provide insight into BP effects upon individual organs or the potential for BP-induced transcriptomic changes. To test the transcriptional impact of ZOL outside of the skeleton, 7 non-skeletal tissues were collected from aged (24mth, F) mice following ZOL treatment (2 months, 125μg/kg or vehicle control) and compared for local alterations in gene signatures (using Nanostring GeoMx platform, Bruker). In each individual tissue, representative ROI across organs were selected using IF-stained (CD45, panCK) sections based on key tissue-specific regions (**Fig.2h** (e.g. heart, liver)). These showed that ZOL treatment triggers a variety of tissue-specific transcriptional changes at non-skeletal sites. These included 96 genes in liver (Top 3: *Scd1, Serpine1, Car3*), 44 genes in heart (Top 3: *Ankrd1, Corin,* and *Ankrd23*), 18 genes in spleen (Top 3: *Fer1l6*, *Ihh,* and *Vmn1r189*), 9 genes in intestine (Top 3: *Alb*, *Apoa1*, *Apoe*), and 7 genes in lung (Top 3: *Gm31714*, *Pgrmc1*, *Sesn3*) (**Fig.2e, 2f**). Simultaneously, 17 genes were commonly modulated in 2 or more organs, of which *Sesn3* was upregulated in heart, spleen and lung while downregulated in intestine. *Acadm*, *Rps3a1*, *Eif1*, *Hadh*, *Ech1*, *Cox6c*, *Acaa2,* and *Acat1* were upregulated in both heart and liver. *Gm31714*, *Pramel25,* and *Gm11554* were both upregulated in spleen and lung while *Chic2* and *Dhrs9* were upregulated in spleen and downregulated in lung. *Cyp2d12* was downregulated in both heart and lung. *Crybb2* was downregulated in both intestine and spleen. *Rpl21* was upregulated in both heart and spleen. Moreover, ageing-related genomic signatures were positively altered in organs from ZOL-treated animals. Based on these results, 32 genes in liver, 2 genes in intestine, 10 genes in the spleen, and 8 genes in the heart were mapped to the HAGR databases (**Fig.2g**). The single cell deconvolution results indicate that the organ-based cell landscape vastly differed between young and aged mice. However, upon ZOL administration, aged mice exhibited a cell distribution shift towards that of younger animals in heart, liver, and lung (**Fig.2i**).

### Bisphosphonate uptake and distribution in non-skeletal cells

Through their binding of calcified surfaces (bone) and internalization by bone-resorbing osteoclasts, the effects of BP have been thought to be primarily restricted to these cells alone. However, other phagocytic cell populations of the bone microenvironment (and beyond) may also have potential for BP internalization^82,83^ with possible effects beyond bone^84^. Having demonstrated systemic proteomic and local genomic effects of BP treatment *in situ*, we sought to validate and visualize the internalization of BPs within a panel of non-skeletal human cell types. Fluorescently labeled RIS (ROX-RIS) and ZOL (FAM-ZOL, ATF-ZOL) were used alongside viral based fluorescent organelle markers (**Fig.3a**). Whilst uptake efficiency varied across cell types, both RIS and ZOL were observed within the majority of cells tested (**Fig.3b**) (0.1µM), with heart, kidney, and liver cells demonstrating the highest BP content, and indicating that the intracellular availability of BPs might vary among diverse tissues. Co-IF imaging with a panel of intracellular organelle markers revealed a preferential localization of RIS and ZOL with lysosomes and endosomes, suggesting a potential role for BPs in the management of intracellular protein quality, quantity, and modification (**Fig.3c and 3d**). These observations not only underscore a broader cellular accessibility of BPs than previously reported, but also hint at potential intracellular targets and pathways that might be influenced.

**Figure 3.**
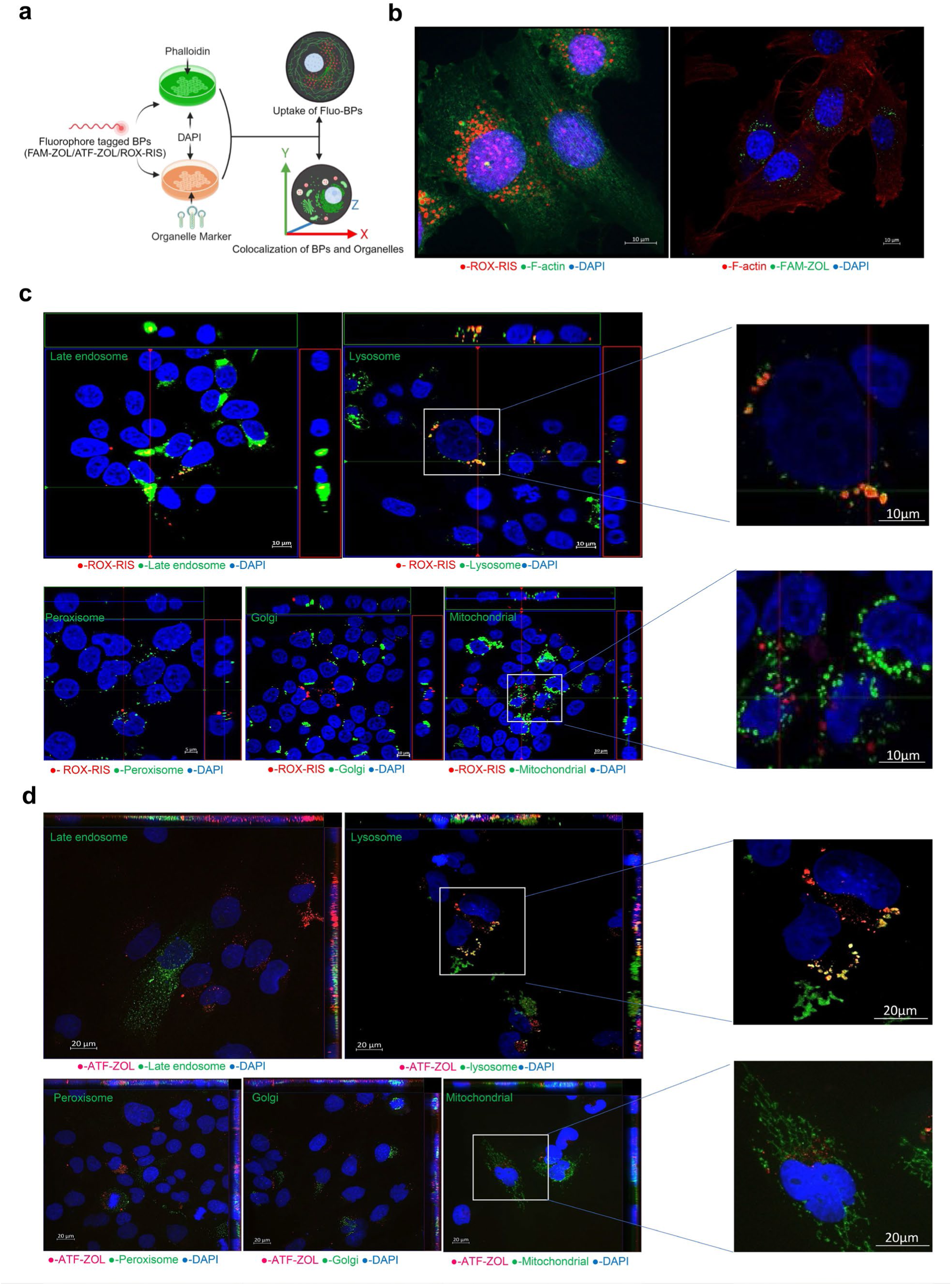
Uptake and subcellular distribution of BPs in non-skeletal cells. **a.** Graphical abstract depicting fluoro-BP distribution/co-localization workflow using ROX-RIS, FAM-ZOL, ATF-ZOL (0.1 μ M), DAPI, Phalloidin (0.1 µg/ml), and organelle-markers of late endosome, lysosome, peroxisome, Golgi apparatus, and mitochondria. Z-stack applied for 3D analysis of confocal microscopy. **b.** Subcellular distribution of ROX-RIS and FAM-ZOL in HUH-7 cells. **c.** Localization of ROX-RIS and organelles in HEK cells, and **d.** ATF-ZOL in AC-16 cells.

### Bisphosphonates induce contrasting dose-dependent effects on cell growth

The internalization of relatively high doses of BPs by osteoclasts during bone resorption (approximately 10μM) leads to apoptosis^85^. A panel of human cells was treated with clinical and novel BPs across multiple doses and time points to explore the full cellular impact of BPs. In considering the functional diversity within the BP family, seven typical compounds were selected representing non-nitrogen (non-N) containing BPs (CLO, ETI), nitrogen (N) containing BPs (ZOL, RIS, ALN), and two non-clinically used nitrogen containing BPs (OX14, IG9402) with varying bone affinity and effects on FPPS. Compounds were tested in cells across representative tissue types, including those linked to beneficial clinical outcomes (lung, heart, blood vessel, kidney, muscle, liver, prostate, immune, and stem cells) (**Fig.4a**). Using live cell imaging (IncuCyte) combined with AI-aided confluence analysis, BP treatment (up to 4 days) at low (0.001μM), mid (0.1μM), and high (10μM) doses revealed contrasting effects across multiple cell types (**Fig.4b, c**). At high BP doses, a significant reduction in growth was induced by the majority of BPs tested, similar to that seen with apoptotic control and in agreement with our understanding of BP action in osteoclasts. However, low BP doses (<0.01μM) were shown to stimulate cell growth. Specifically, low dose treatment with ZOL significantly stimulated proliferation in cardiomyocyte (28%), cardiovascular endothelia (9.4%), renal epithelia (20.5%), hepatocytes (22%), myoblasts (8.7%), and monocytes (21.1%). A similar significant increase was seen with RIS and the non-N BP CLO (**Fig.4d, e**). These data suggest that BPs might act upon a variety of cell types, with the potential to confer beneficial effects at lower doses. Importantly, as N- and non-N-containing BPs function through distinct cellular mechanisms, the similar increase in cell growth stimulated by most BPs suggests combined or novel mechanisms might mediate this response.

**Figure 4.**
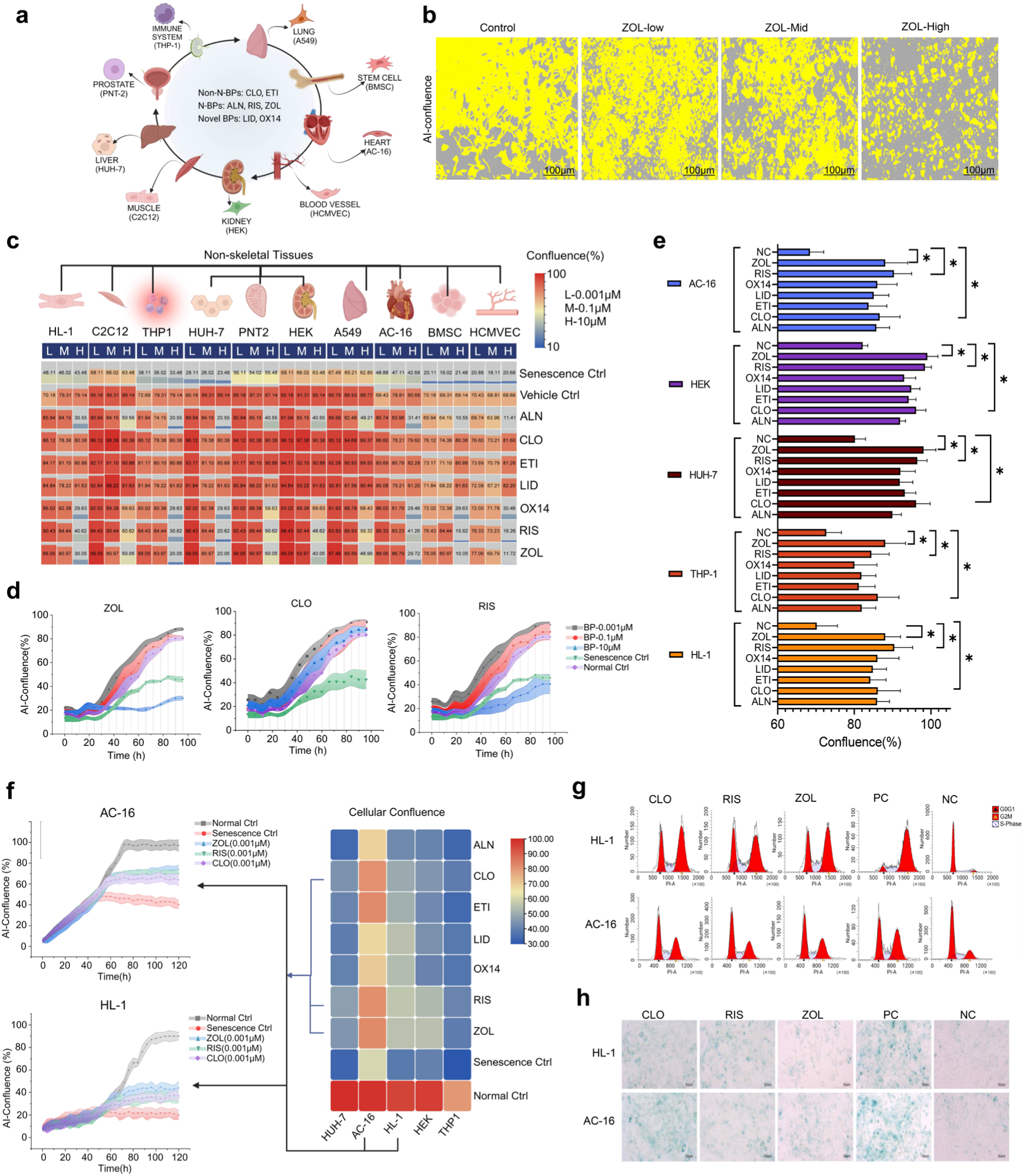
BPs alter non-skeletal cell biology and growth. **a.** Screening protocol for BPs and cell panel. **b**. Representative confluence analysis (HUH-7) from **c.** Endpoint confluence ratio of non-skeletal cell panel following ZOL treatment (L=0.001, M=0.1, H=10μΜ, 96 hrs). **d.** Time course effects of ZOL, CLO, RIS on cell growth (representative HUH-7). **e.** Low dose BP related growth (AC-16, HL-1, HEK, HUH-7, THP-1), and **f.** following MMC-induced senescence induction (time course-CLO, RIS, ZOL and end point confluence-heatmap). **g.** BP-pretreatment on senescence markers (γ-H2AX, LaminB1, P16, IL-6, TNF-α and HMBG1) following MMC treatment (AC-16, PC: 0.05μg/ml MMC; NC: vehicle), **h.** cell cycle (PI-labelled), and **i.** SA-β-Gal staining (x200). MMC treatment=senescence control. * p<0.05 vs normal control (NC). Data presented as mean±SD, n=3.

### Bisphosphonates protect against DNA damage-related senescence

Expanding upon our observations on ageing and senescence from BP-treated human and murine studies, we tested whether BP pre-treatment might reduce senescence induction in our panel of non-skeletal cells. Exposure of HUH-7, AC-16, HL-1, HEK and THP-1 cells to low dose BP (0.001μM) prior to induction of DNA damage-triggered senescence, protected against a reduction in confluence, with CLO, RIS, ETI, and ZOL demonstrating the highest degree of protection in cardiomyoblast cells (**Fig.4f**). Furthermore, significant changes in protein markers were observed reflecting reduced senescence, including expression of γ-H2AX, LaminB1, P16, IL6, and TNF-α, cell cycle suspension (**Fig.4g**), and SA β-Gal staining (**Fig.4h**), following pre-treatment with ZOL, CLO, or RIS.

### Global profiling of Zoledronate-binding proteins

Based on previous evidence indicating ZOL-treatment favorably alters the plasma proteome of osteopenic patients, triggers beneficial genomic changes at multiple organ sites *in vivo*, and promotes cell growth at low doses in multiple human cell types *in vitro* including human cardiomyoblast cells, we prioritized ZOL for 2D thermal profiling to investigate its proteome-wide target spectrum. This method measures protein stabilization at target saturating compound concentrations upon application of a heat gradient. First, we confirmed by western blot that the known ZOL target farnesyl pyrophosphate synthase (FPPS) was stabilized upon ZOL treatment over increasing temperatures (37-78℃) in intact A549, HEK293, AC-16, and HUH-7 cells. AC-16 cardiomyoblasts showed the most significant thermal stabilization effect across all four cell lines (ΔT_m_ =17.38℃) (**Fig.5a**). Protein lysates of intact AC-16 cardiomyoblasts collected at increasing temperatures with different concentrations of ZOL (0-20μM) were further analyzed using mass spectrometry and TP-MAP^86^ to identify proteins that might be positively or negatively (destabilized) altered by ZOL binding. Reassuringly, we were able to identify the cognate target FPPS, as the highest scored protein across 8397 identified proteins, following ZOL treatment (**Fig.5b**). In addition, several other proteins so far not associated with BPs, were also significantly stabilized following ZOL treatment, with direct target engagement confirmed by western blot (**Fig.5c**) for PHB2, ASAH1, and FOSL1 (ΔT_m_ PHB2=12.15 ℃, ΔT_m_ ASAH1=12.13 ℃, ΔT_m_ FOSL1=2.09 ℃). PHB2 is a multifunctional protein involved in various cellular processes such as mitochondrial function, cell cycle regulation, and apoptosis^87^. ASAH1 encodes an enzyme involved in the metabolism of ceramides, which plays a role in various cellular processes, including cell signaling and apoptosis^88^, whilst FOSL1 is a crucial component of the AP-1 transcription factor complex playing a key role in regulating cell proliferation, differentiation, and tumorigenesis^89^.

**Figure 5.**
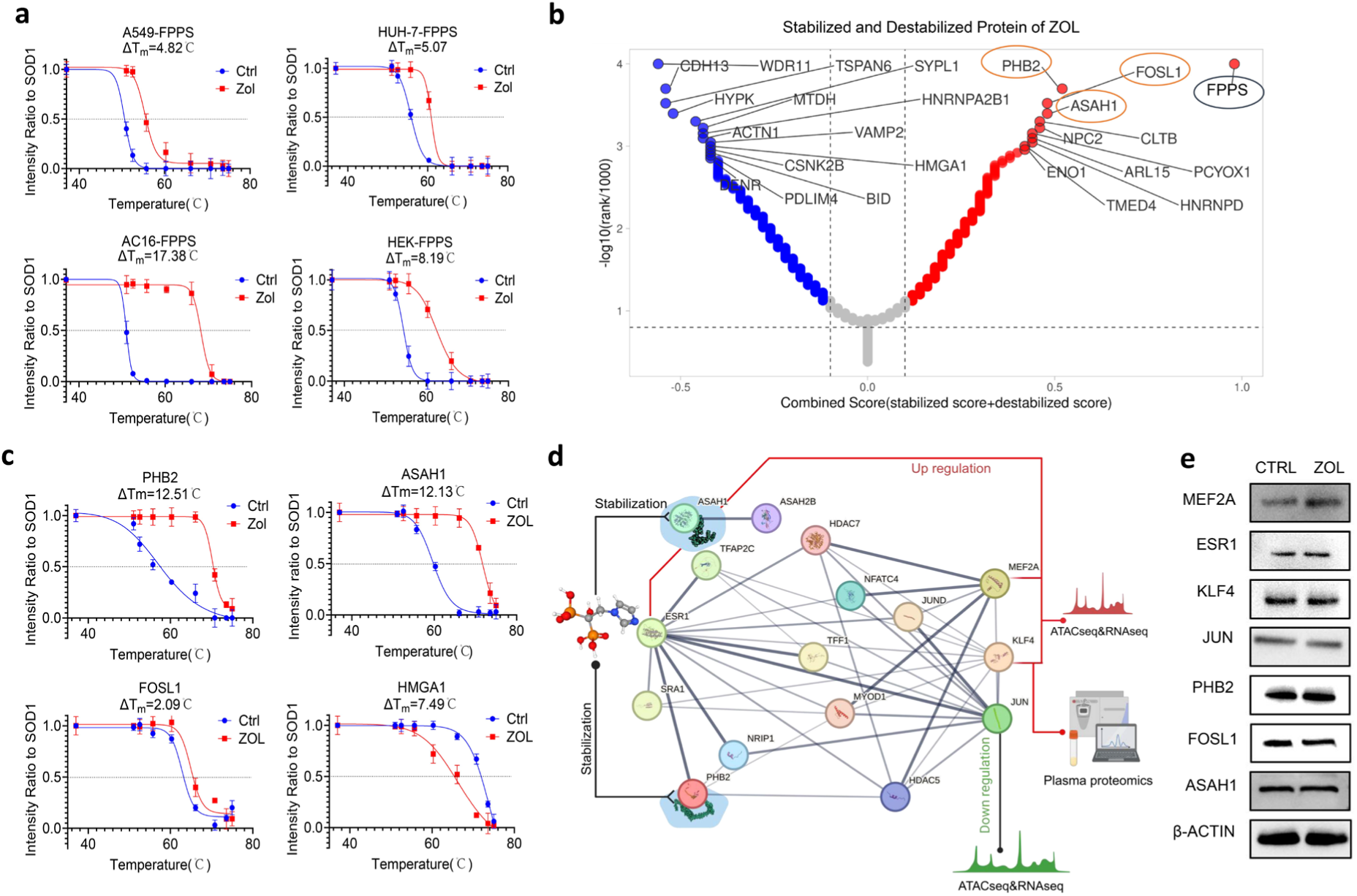
Zoledronate binding targets. **a.** FPPS thermal stability profiles following ZOL/vehicle ctrl treatment (A549, HEK293, AC16, HUH-7) relative to SOD1 (internal control). **b.** ZOL stabilized/destabilized proteins (AC-16) following 2D thermal profiling. **c.** CETSA**-**western blot validation of thermal profiling hits (ASAH1, PHB2, FOSL1, HMGA1). **d.** Cross-omic analysis following ATAC-seq and RNA-seq of ZOL-treated (0.001μM) AC-16 cells, downstream of prioritized CETSA targets (PHB2 and ASAH1). Further mapped to human proteome analysis following ZOL treatment (using STRING). **e.** Validation of ZOL-induced protein changes (MEF2A, ESR1, KLF4, C-JUN, PHB2, FOSL1, ASAH1). Data presented as mean±SD, *p<0.05 vs control (Ctrl), n=3.

### Zoledronate triggers transcriptomic changes in cardiomyoblasts

To systematically explore the regulatory network associated with ZOL-stabilized protein targets, RNA- and ATAC-seq analysis were performed on AC-16 cardiomyoblasts with/without ZOL treatment to identify changes in gene transcription and chromatin accessibility, respectively. Cells were sequenced using NextSeq 500 following ZOL treatment (0.001 μM, 4 days) or vehicle control. An expanded protein-protein interaction network (PPI), originating from PHB2 and ASAH1, was constructed using the STRING database. All factors related to these targets were mapped to our RNA-, ATAC-seq, and SomaScan proteomic datasets to identify genes/proteins common to all analyses and altered in human systems (*in vitro* and *in vivo*) following exposure to ZOL. This integrated analysis indicated four potential regulators (ESR1, MEF2A, JUN, KLF4) downstream of identified protein targets of ZOL (**Fig.5d**). Western blot analysis confirmed sustained translational impact of MEF2A only, showing increased protein expression following ZOL treatment (**Fig.5e**). The MEF2A transcription factor (myocyte enhancer factor 2a) plays a central role in many biological processes, particularly in muscle development and cardiovascular biology^90^. Moreover, MEF2A may suppress senescence in vascular endothelial cells^91,92^. Using public databases specific to transcription targets (CHEA, TTRUST, CISTRONE, ENCODE) a pool of MEF2A targets in human tissues was developed (**Fig.6a**). A total of 5218 human gene targets of MEF2A were identified, 2208 of which mapped to at least one of our ATAC-, RNA-seq, or SomaScan proteomic datasets (**Fig.6b**). Of those gene targets, 154 were identified in both ATACseq and RNAseq while 40 were identified in all three omic datasets (**Fig.6b**). The regulatory impact of MEF2A is indicated along with alterations in 16 targets in one or more established Ageing databases (GenAge, LongevityAge, CellAge) (**Fig.6c**). These were HMBGB1, HSP90AA1, JUN, MAPK1, PIK3R1, TFRC, PURB, PDLIM1, NT5E, LYN, HSPH1, HMGA1, FN1, CLIC4, TGFB1, and VCAM1. For all mapped MEF2A targets, a pathway enrichment analysis was also conducted to illustrate potential biological processes influenced by ZOL-activated MEF2A. Of the 17 functional groups indicated, heart disease/angiogenesis, cell cycle, musculoskeletal morphogenesis, hearing/ear morphology, inflammation, infection/defense reaction demonstrated the highest degree of significance (**Fig.6h**). Furthermore, previous RNA-seq analysis following knockdown of MEF2A in cardiomyoblasts^91^ was cross-referenced with our ZOL-treated datasets (and similarly using the KnockTF 2.0 database, which profiles gene expression following transcription factor knockdown/knockout across multiple species) (**Fig.6d and 6e**). We identified 15 common targets showing inverse relationships between ZOL-stimulated MEF2A increase in our human proteomic, RNA-, ATAC-seq and MS analysis, and MEF2A knockdown (**Fig.6f**). These were SULF2, NMT2, TACSTD2, FAHD1, ARB1, RSPO3, SULT1B1, RNASE1, SULT1E1, SAT1, LGALS3BP, CPA4, CBS, GSN, PIK3R1 (**Fig.6g**). A heatmap was generated to illustrate the detailed regulatory information and mapping results from the HAGR^93^ ageing databases. From these, TACSTD2, GSN, and PIK3R1 were indicated across three omics-mapped target pools and two omics-mapped knockout pools (**Fig.6c, g**), highlighting a previously unreported role in BP regulation and as potential ageing-related factors. This work indicates BPs function in a dose dependent manner where low doses offer protection against the onset of cellular senescence outside of bone, and which may be mediated through the binding of novel targets (PHB2, ASAH1), activation of MEF2A, and regulation of downstream targets including PURB, MAPK1, CBS, TGFB, HSP90AA1, and HMGA1 (**Fig.6i**).

**Figure 6.**
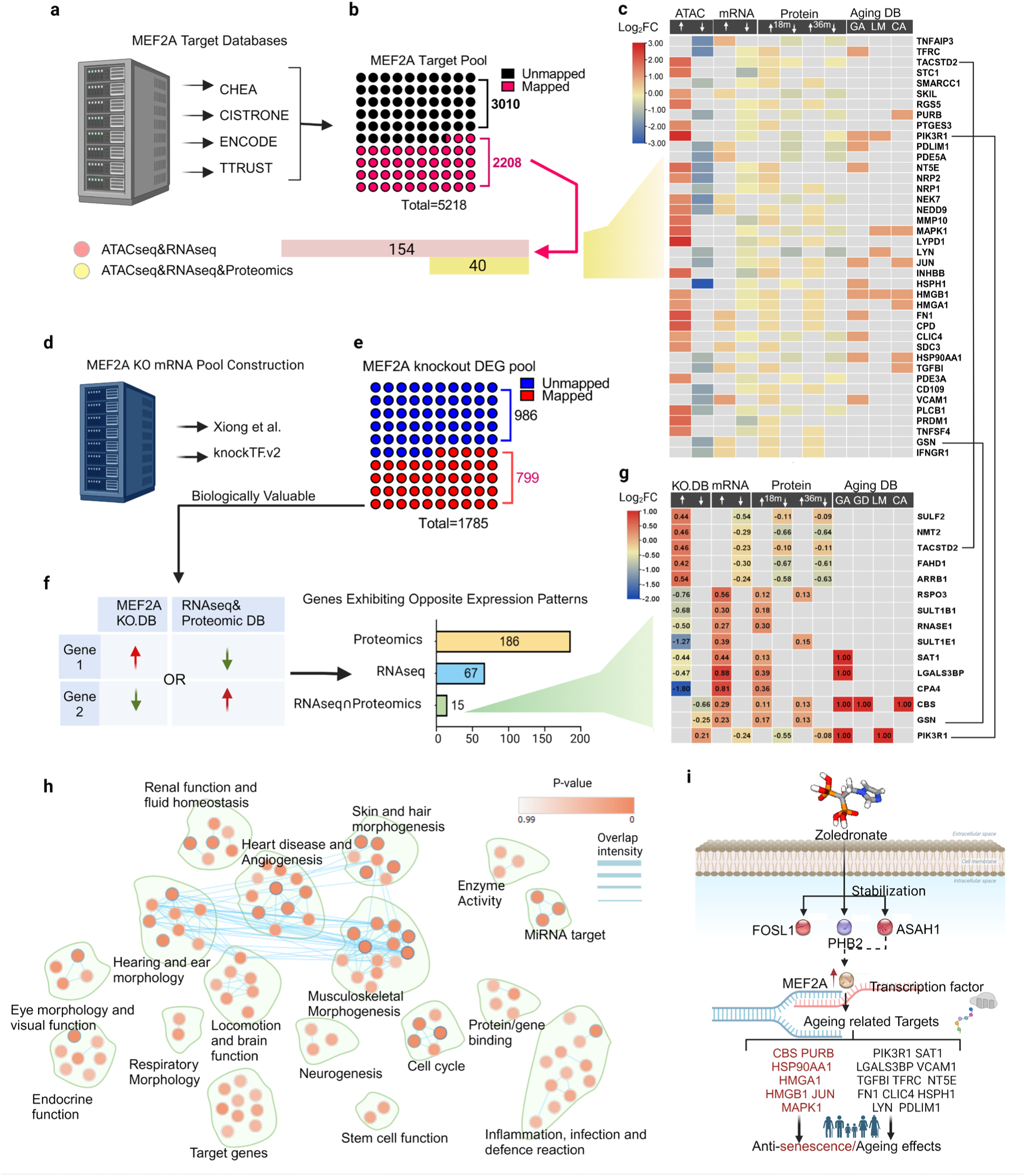
Cross-Omic functional analysis reveals ZOL-MEF2A axis. **a.** Human tissue-derived gene targets of MEF2A from TF target databases (CHEA, TTRUST, CISTRONE, ENCODE) revealing **b,** MEF2A targets mapped to ATAC-seq, RNA-seq, and human proteome datasets, and **c**. HAGR (GenAge (GA), GenDR (GD), LongevityMap (LM), and CellAge (CA)). **d**, MEF2A knockout pool targets revealing **e**, DEGs following MEF2A knockout **f,** mapped to ATAC-seq, RNA-seq, and human proteome datasets, and **g,** HAGR databases. **h**. Pathway enrichment analysis of mapped ZOL-MEF2A axis targets (GSEA) as bio-functional categories. **i**. Graphic summary ZOL-MEF2A axis.

## Discussion

While it remains clear that the calcium-binding property of BPs, along with their capacity to bind and inhibit FPPS in bone-resorbing osteoclasts^94^ constitutes the major effect and mechanism of this class of drugs in humans, this study highlights key areas where and how BP treatment elicits new effects. BP use varies across the globe, with trials primarily focused upon white female populations in developed regions such as the United States, United Kingdom, and New Zealand. Given the genetic and environmental variations influenced by gender, race, and geography, the impact of BPs on disease progression and mortality reduction could significantly differ across different demographics. Improved information on regions, races, and particularly male subjects, will be needed to form a complete picture of BP impact. However, a strong and clear pattern indicating significant benefit of BP treatment across multiple disorders has emerged, and where a single mechanism of action would be unlikely. In beginning to explore this polypharmacology of BPs, we have uncovered unique binding partners and downstream regulators which may collectively impact lifelong human health and prevent disease.

Treatment with ZOL in murine and human studies triggered significant genomic and proteomic changes respectively, in multiple tissues. This included alterations in immune cell regulation and changes to inflammatory factor profiles. In support of these findings, BPs have been shown to activate γΔT-cell populations^95^. This phenomenon may account for the acute phase response triggered by some BPs, that leads to flu-like symptoms following start of treatment^96,97^. Patients treated with ZOL also upregulated proteins controlling autophagy, a natural process of waste disposal and recycling, that decreases with ageing, preventing the production of defective proteins. Autophagy is dependent upon effective endosomal/lysosomal activity to capture, internalize, and degrade defective proteins within an autophagosome^98^. Our combined fluoro-labeled BPs and proteomic analysis tracked internalized BPs to these intracellular loci and confirmed elevated circulating levels of protein markers of lysosomal activity. In a similar approach to that adopted in our study, Yu, Surface, and colleagues identified new BP transporter molecules using CRISPR-mediated genomic analysis following ZOL-treatment. Through a SLC37A3-ATRAID complex, bound BPs transported through the cytosol are released at lysosomal sites^99^ supporting a role for BPs in ageing-linked autophagy control and highlighting the possibility of further BP binding targets.

BPs have been reported to promote tumour cell apoptosis through activation of autophagic cell death in a range of cancers including prostate, breast, cervical, colon, and osteosarcoma. This is suggested to occur via reduced activity of the mevalonate pathway enzyme, geranylgeranyl diphosphate (GGDP). However, this could not be rescued through re-addition of mevalonate pathway metabolites or use of other pathway inhibitors, suggesting BP action is only partly mediated by mevalonate pathway inhibition and where other mechanisms (e.g. inhibition of mTOR signaling) will play a role^100–105^. Modification of mTOR signaling by BPs has also been linked to survival in mesenchymal stem cells, where ZOL conferred protective effects against DNA-damage^106^, and improved survival in Drosophila^107^. This might also account for rejuvenating effects seen in our animal studies where ZOL-treatment in aged mice induced a tissue-specific (heart, liver, lung) local genomic change, reflecting that of young animals.

A strong and significant change in cellular senescence may also have occurred following BP treatment, both in senescence induction in these tissues, or in the targeting and removal of senescent cells following ZOL treatment. Non-proliferative senescent cells accumulate in tissues with increasing age. These cells corrupt the local environment by secreting a toxic cocktail including inflammatory factors and matrix-degrading enzymes^108^, whilst removal of senescent cells improves tissue function^109^. Collectively, our work shows BP-treatment reduces formation of senescent cells and products (SASP)^110^, where alterations in senescence-associated gene signatures across local tissues is supported by changes in downstream protein production in humans treated with ZOL and in individual cell-specific cultures, such as BP-induced changes across multiple inflammatory markers.

Furthermore, we have revealed how new targets of BPs may confer such beneficial effects. Integrated analysis of our cellular and omics studies has revealed a unique axis downstream of new BP-binding targets, where PHB2-MEF2A activation offers protection against the induction of senescence, and potential onset of an ageing phenotype. The prohibitin family (PHB) are highly conserved, widely expressed pleiotropic proteins implicated in a variety of cellular functions (proliferation, apoptosis, transcription, tumor suppression). Importantly, preservation of PHB2 activity is linked to increased lifespan brought about by improved mitochondrial metabolism^111,112^ and similar to BPs, protection against cardiovascular disease and neurodegeneration^113–115^. Similarly, MEF2A expression declines in ageing tissues, whilst MEF2A activation may improve skeletal muscle function in aged animals, cognitive decline in neurodegenerative models, and endothelial cell function in cardiovascular disease^92,116–118^. Furthermore, mutations in MEF2A have been reported in families with CVD^119–121^. In support of our findings, MEF2A deletion led to increased senescence in coronary artery endothelial cells^91^ suggesting that MEF2A activation following BP treatment is protective in this system. Our research presents a comprehensive analysis of BP effects beyond their traditional use in preserving bone health, where their impact on serum proteomics, cellular ageing, and immune cell rejuvenation offers new therapeutic possibilities. These findings expand our current understanding of the operational mechanisms of BPs within human cells, and impact upon local tissues *in vivo* and systemic effects in humans. These suggest both FPPS-dependent and new independent mechanisms through which BPs might confer beneficial effects in ageing-related disorders, highlighting exciting opportunities for BPs as new gerotherapeutic drugs.

## Acknowledgements

This work was supported by funding from the UKSPINE Knowledge Exchange Flagship Fund (JE), the US National Institutes of Health grants P01 AG062413 and R01 AG086085 (SK), and JLK supported by R37AG13925, R33AG61456, R01AG072301, R37AG13925, and the Hevolution Foundation (HF-GRO-23-1199148-3), the Connor Fund, Robert J. and Theresa W. Ryan, and the Noaber Foundation. In addition to indicated authors/collaborators, we are grateful to James Dunford, and Martin Philpott and Adam Cribbs for knowledge and assistance with bisphosphonate compounds, and ATAC-seq and RNA-seq analysis, respectively.

## Contributions

JL, HZ, and HK performed cellular studies. SR, JL performed bioinformatic analysis on all data generated. Human ZOL-treated samples provided by IR, AH. HZ, JL, KH, LB, IV and BK were responsible for 2D thermal profiling and CETSA analysis. ER, GR, HE, BG provided expertise in bisphosphonate chemistry including fluorophore tagged BPs and distribution. SK provided ZOL-treated ageing mouse samples. EP, LF, TJA, GM performed spatial transcriptomics analysis. JK, CB, GR, JE contributed to study design, direction, and interpretation of data. JE oversaw study design, management, interpretation of data. JL, JE prepared the manuscript with input from all authors.

